# Mammalian Cell Culture and Transfection in a Low-Cost, CO_2_-Independent Environment

**DOI:** 10.1101/2025.10.14.682406

**Authors:** Antonio Lamb, Aron Gagliardi, Dominique Lorgé, Lucia Carillo

## Abstract

Traditional mammalian cell culture requires CO_2_ incubation to maintain physiological pH, creating infrastructure costs that limit accessibility. We demonstrate successful HEK293T/17 culture using HEPES-buffered CO_2_-independent medium in a repurposed consumer-grade egg incubator, reducing incubator equipment costs by >98% while maintaining viability through 10+ passages. To assess feasibility for recombinant protein production, we performed plasmid transfection with pcDNA3.1(+)-EGFP via electroporation. Fluorescence microscopy confirmed robust EGFP expression within 48 hours. These results demonstrate that mammalian cell culture and genetic engineering for recombinant protein production can be achieved using cost-effective, widely available equipment rather than conventional high-cost laboratory infrastructure.

## Description

Traditional mammalian cell culture relies on CO_2_ incubation systems to maintain physiological pH through bicarbonate buffering. The CO_2_/bicarbonate buffering system maintains pH by equilibrating gaseous CO_2_ with dissolved bicarbonate in the culture medium, requiring precise control of atmospheric CO_2_ concentration, typically at 5-10%. While effective, this approach creates significant infrastructure costs that limit accessibility for resource-constrained laboratories, educational institutions, and researchers in developing regions. Standard CO_2_ incubators cost several thousand Swiss Francs, require regular maintenance, and depend on continuous CO_2_ gas supply. These barriers substantially limit the democratization of cell culture research and restrict opportunities for small-scale laboratories to engage in mammalian cell culture.

Alternative buffering strategies using zwitterionic buffers such as HEPES (4-(2-hydroxyethyl)-1-piperazineethanesulfonic acid) offer CO_2_-independent pH maintenance (Good et al., 1966). HEPES maintains stable pH in the physiological range (6.8-8.2) without requiring CO_2_ supplementation (Michl et al., 2019). Despite the availability of CO_2_-independent media formulations, CO_2_ incubation remains entrenched in cell culture practice, largely due to historical precedent and the perception that specialized equipment is necessary for all mammalian cell culture applications. We hypothesized that combining HEPES-buffered CO_2_-independent medium with low-cost temperature and humidity control could provide a viable alternative to traditional CO_2_ incubation systems for basic cell culture applications.

To test this hypothesis, we selected HEK293T/17 cells, a widely used human embryonic kidney cell line commonly used for protein expression, transfection studies, and viral vector production (Lomba et al., 2021). We repurposed a consumer-grade egg incubator (NEWFUN, Amazon.de Cat# BOF4WY21YV, 82.00 CHF) to provide temperature and humidity control without CO_2_ supplementation. The egg incubator was selected based on its ability to maintain stable temperature (37°C) and high humidity (85%), which are critical parameters for mammalian cell culture independent of CO_2_ control. We formulated culture medium using 1:1 DMEM:F12 supplemented with 10% fetal bovine serum (FBS), antibiotics (100 I.U./mL penicillin, 100 µg/mL streptomycin), and 15 mM HEPES buffer adjusted to pH 7.4.

Cells were revived from cryopreservation and cultured in this system, with media changes performed every 2-3 days when the pH dropped to 7.0, based on phenol red pH indicator color. Phenol red shifts from orange-red (pH 7.4) to yellow (pH <7.0) as culture medium acidifies due to cellular metabolism, providing a visual cue for media replacement (Using Phenol Red to Assess pH in Tissue Culture Media, 2018). Environmental monitoring over more than 40 hours using a temperature and humidity data logger (Tzone Model #T119) confirmed that the egg incubator maintained stable conditions at 37.0 ± 0.5°C and 85 ± 3% relative humidity (Figure 1D). These parameters fall within the acceptable range for mammalian cell culture and are comparable to those achieved in standard laboratory incubators.

**Figure 1.**
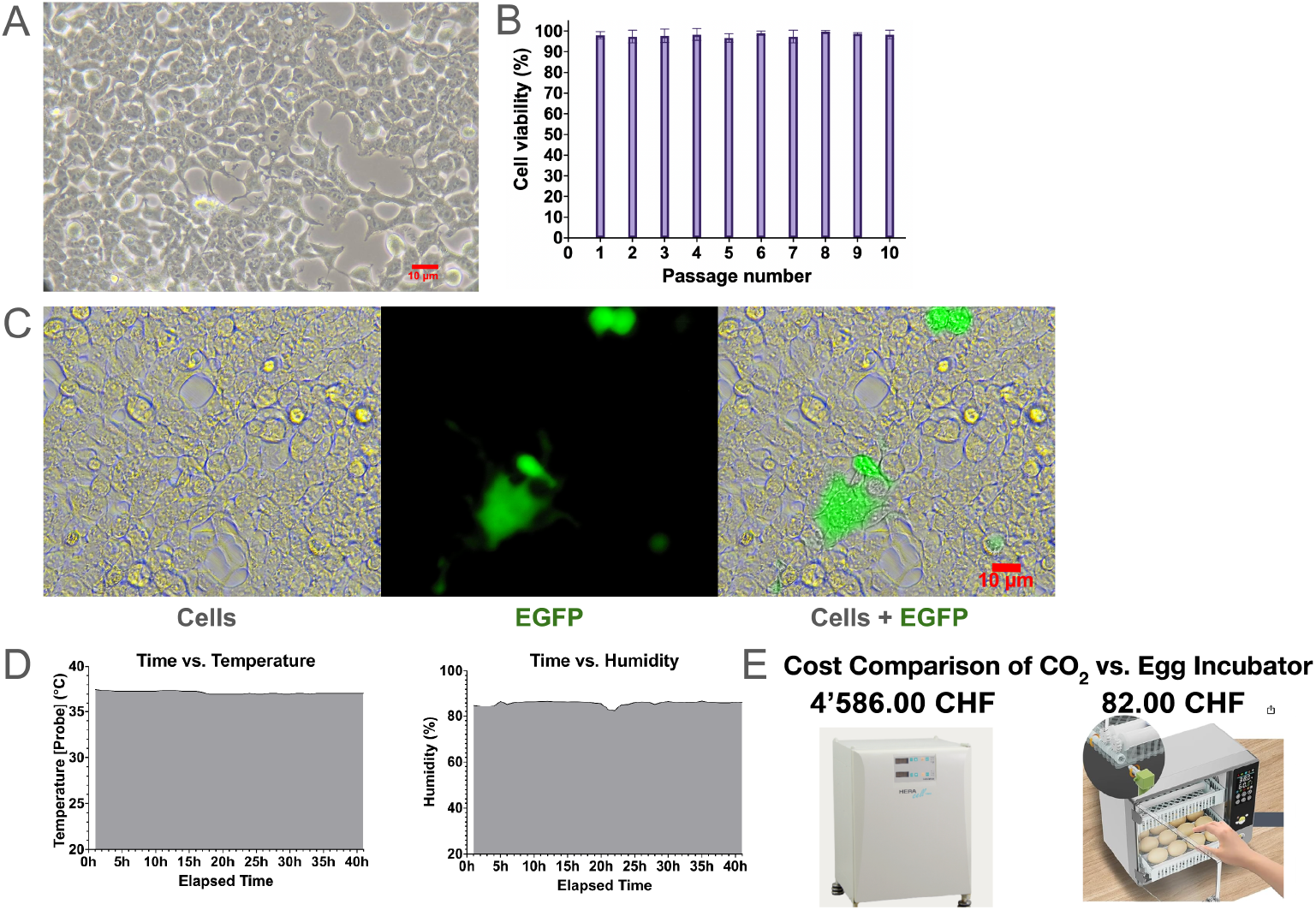
HEK293T/17 cell culture and transfection in CO_2_-independent conditions.: **(A)** Representative 20X phase contrast microscopy image showing healthy HEK293T/17 cell morphology in 1:1 DMEM:F12 + 10% FBS + 100 I.U./mL penicillin + 100 µg/mL streptomycin + 15mM HEPES at passage 4, cultured in a consumer-grade egg incubator at 37°C in ambient air. Scale bar = 10 μm. **(B)** Cell viability quantified by trypan blue exclusion assay with consistent viability (>95%) maintained across 10 passages without CO_2_ supplementation. Error bars represent standard deviation (n=3 technical replicates per passage). **(C)** Transfection of HEK293T/17 cells with pcDNA3.1(+)-EGFP plasmid via electroporation using HEPES-buffered electroporation medium. Left panel: brightfield image showing cell morphology 72 hours post-transfection. Middle panel: fluorescence imaging (Ex/Em 490/525 nm) of EGFP expression. Right panel: merged brightfield and fluorescence channels. Scale bar = 10 μm. **(D)** Environmental monitoring of egg incubator configured to 37°C (left) and 85% relative humidity (right) over 40-hour period. **(E)** Cost comparison between traditional CO_2_ incubator (4,586 CHF) from Thermo Fisher and consumer-grade egg incubator (82 CHF).

Cell viability was assessed across 10 consecutive passages using trypan blue exclusion assays performed in triplicate. At each passage, cells were harvested at greater than 80% confluence using standard trypsinization (0.25% Trypsin-EDTA, 300×*g* centrifugation for 3 minutes) and viable cell counts were determined using a hemocytometer. Cell viability remained consistently above 95% across all 10 passages, with consistent cell density exceeding 2.78×10^6^ cells/mL achieved at confluence (Figure 1B). Microscopy revealed healthy cell morphology characteristic of HEK293T/17 cells, with no evidence of stress-related morphological changes such as cytoplasmic granulation or detachment (Figure 1A). These results demonstrate that HEPES-buffered medium provides robust pH stability sufficient to maintain HEK293T/17 cell viability over extended culture periods without CO_2_ supplementation.

To further validate the functionality of cells cultured under CO_2_-independent conditions, we performed transfection experiments using the pcDNA3.1(+)-eGFP plasmid encoding enhanced green fluorescent protein. Transfection capability is a critical functional assay for HEK293T/17 cells, as this cell line is widely used for recombinant protein expression. We used electroporation using a Bio-Rad Gene Pulser 1 system with cells resuspended at 5×10^6^ cells/mL in electroporation buffer (10 mM HEPES pH 7.4, 285 mM sucrose, 0.9 mM MgSO_4_·7H_2_O, 1.3 mM KCl). Electroporation parameters were set at 200 V, 25 µF capacitance, and 200Ω resistance in 0.2 cm cuvettes, with 2.66 µg plasmid DNA added to 160 µL cell suspension. Following electroporation, cells were incubated for 10 minutes at room temperature before being transferred to the egg incubator at 37°C and 85% relative humidity.

Fluorescence microscopy performed 48 and 72 hours post-electroporation revealed robust EGFP expression in transfected cells. Time constants recorded during electroporation ranged from 2.1-2.2 ms, indicating successful membrane permeabilization and plasmid delivery. Fluorescence imaging using an AE-IM400 inverted microscope with appropriate excitation (490 nm) and emission (525 nm) filters confirmed functional EGFP expression (Figure 1C). The successful expression of recombinant protein demonstrates that cells cultured without CO_2_ supplementation retain full functionality for molecular biology applications, including transfection and protein expression.

Additionally, we established successful cryopreservation protocols compatible with this culture system. Cells were frozen at approximately 1×10^6^ cells/mL in CM-1 freezing medium (Cytion Cat#800100), cooled gradually at 4°C for 4 hours in a cell freezing container before transfer to −70°C storage. After 30 days of cryostorage, cells were rapidly thawed at 37°C, washed with DMEM, and re-cultured in supplemented 1:1 DMEM:F12 medium in the egg incubator. Revival was successful with cells reaching greater than 80% confluence.

The economic advantage of this approach is substantial. The egg incubator costs 82.00 CHF (Amazon.de Item #B0F4WY21YV) compared to 4,586.00 CHF for a standard CO_2_ incubator (Thermo Fisher 11626250, refurbished), representing a greater than 98% cost reduction in initial equipment investment (Figure 1E). When considering the elimination of ongoing CO_2_ gas supply costs, the savings compound even more over time. The reduced cost makes mammalian cell culture accessible to resource-constrained settings, including teaching laboratories, small research groups, and institutions in low- and middle-income countries.

There are several important limitations we identified while culturing HEK293T/17 cells in ambient environments. HEPES buffering has a finite buffering capacity, requiring more frequent media changes (every 2-3 days) compared to CO_2_-buffered systems where media may last longer. The phenol red pH indicator serves as a useful visual guide, but regular monitoring is still necessary to prevent rapid and excessive pH swings. While our results demonstrate robust performance with HEK293T/17 cells, other cell lines with different metabolic rates or pH sensitivities may require optimization of HEPES concentration or media change frequency. The egg incubator lacks the sophisticated digital control and uniformity of commercial cell culture incubators, though our monitoring data indicated that the device was more than sufficient for our applications. Finally, while we demonstrated successful transfection and protein expression, more demanding applications such as long-term differentiation protocols or primary cell culture may require further optimization.

In summary, we demonstrate a practical, low-cost alternative to traditional CO_2_-dependent mammalian cell culture using HEPES-buffered medium and repurposed consumer equipment. Cells remain viable for transfection and recombinant protein expression in ambient atmospheric conditions, reducing incubator infrastructure costs by more than 98%. Our assessment of these protocols gives the general public more equitable access to biotechnology and drug development.

## Methods

### Culture Setup and Maintenance

Cells were revived from cryopreservation and cultured in supplemented 1:1 DMEM:F12 medium at 37°C and 85% humidity inside the egg incubator. Media changes were performed every 2-3 days, or when the phenol red pH indicator showed a pH below 7.0. Temperature and humidity were monitored every hour using a data logger for more than 40 hours to verify stable environmental conditions.

### Cell Passaging and Viability Assessment

Cells were passaged at greater than 80% confluence using standard trypsinization with 0.25% Trypsin-EDTA, followed by centrifugation at 300×*g* for 3 minutes. Cell counting and viability assessment were performed using a hemocytometer with 1:1 dilution of trypan blue, with measurements performed in triplicate. Cells were seeded into T25 flasks at 10,000 cells/cm^2^ and viability was assessed across 10 consecutive passages.

### Electroporation and Transfection

Cells were adjusted to a final concentration of 5×10^6^ cells/mL in electroporation buffer. The pcDNA3.1(+)-eGFP plasmid (2.66 µg total) was added to 160 µL cell suspension and electroporated in a 0.2 cm cuvette using the following settings: 200 V, 25 µF, 200Ω resistance. Following electroporation, cells were incubated for 10 minutes at room temperature, then transferred to the egg incubator at 37°C and 85% relative humidity for 72 hours.

### Fluorescence Microscopy

Image acquisition was performed 72 hours post-electroporation using an AE-IM400 inverted fluorescence microscope with a 250 ms exposure time. Excitation and emission wavelengths were set to 490 nm and 525 nm, respectively.

### Cryopreservation and Revival

Trypsinized HEK293T/17 cells were resuspended in CM-1 freezing medium at 1×10^6^ cells/mL, placed in a cell freezing container, cooled at 4°C for 4 hours, then transferred to −70°C for 30 days. Frozen vials were rapidly thawed at 37°C, washed with DMEM, then seeded into fresh T25 flasks containing supplemented 1:1 DMEM:F12 medium and grown to greater than 80% confluence.

### Reagents

| Cell Line | Species | Genotype/Description | Available from |
| --- | --- | --- | --- |
| HEK293T/17 | *Homo sapiens* | Human embryonic kidney cell line expressing SV40 large T-antigen | Cytion Cat#305117 |

| Plasmid | Genotype/Description | Available from |
| --- | --- | --- |
| pcDNA3.1(+)-eGFP | Mammalian expression vector containing enhanced green fluorescent protein (eGFP) coding sequence under CMV promoter; ampicillin and neomycin resistance | NovoPro Labs<br>Cat#V012862 |

### Cell Culture Media and Buffers

- 1:1 DMEM:F12 supplemented with 10% FBS, 100 I.U./mL penicillin, 100 µg/mL streptomycin, and 15 mM HEPES (pH 7.4)
- Electroporation buffer: 10 mM HEPES (pH 7.4), 285 mM sucrose, 0.9 mM MgSO_4_·7H_2_O, and 1.3 mM KCl
- CM-1 Freeze medium (Cytion Cat#800100)
- 0.25% Trypsin-EDTA
- Trypan blue

### Equipment

- Egg incubator: NEWFUN (Amazon.de Model# BOF4WY21YV), 82.00 CHF
- Electroporation system: Bio-Rad Gene Pulser 1 (Model #1652076), 250.00 CHF
- Temperature/humidity logger: Tzone Model #T119, 40.00 CHF
- -80C Freezer: Alibaba Model# DW-4525, 500.00 CHF
- Inverted fluorescence microscope: AE-IM400, FPlanFL PHP 20X/0.65 objective, Ex/Em 490/525 nm, 2100.00 CHF

## Acknowledgements

We thank the **Swiss Federal Coordination Centre for Biotechnology** for providing guidance and support to ensure that our research activities complied with applicable biosafety regulations and institutional guidelines for cell culture research.

## Funding

This research received no external funding, and was purposely conducted in a low-cost environment to demonstrate that human cell culture and genetic engineering is possible outside of traditional research laboratories. Supported by None to.

## Author Contributions

Antonio Lamb: conceptualization. Aron Gagliardi: methodology. Dominique Lorgé: project administration. Lucia Carillo: writing - review editing.

## Reviewed By

### History: Received

October 13, 2025

### Copyright

© 2025 by the authors. This is an open-access article distributed under the terms of the Creative Commons Attribution 4.0 International (CC BY 4.0) License, which permits unrestricted use, distribution, and reproduction in any medium, provided the original author and source are credited.

### Citation

Lamb A, Gagliardi A, Lorgé D, Carillo L. 2025. Mammalian Cell Culture and Transfection in a Low-Cost, CO_2_-Independent Environment. microPublication Biology. 10.17912/micropub.biology.001899

## Notes

### Competing Interest Statement

The authors have declared no competing interest.

## References

Good NE, Winget GD, Winter W, Connolly TN, Izawa S, Singh RMM. 1966. Hydrogen Ion Buffers for Biological Research*. Biochemistry 5: 467–477. DOI: 10.1021/bi00866a011

Lomba ALO, Tirapelle MC, Biaggio RT, Abreu-Neto MrS, Covas DT, Picanço-Castro V, Swiech K, Mizukami A. 2021. Serum-Free Suspension Adaptation of HEK-293T Cells: Basis for Large-Scale Biopharmaceutical Production. Brazilian Archives of Biology and Technology 64: 10.1590/1678-4324-2021200817. DOI: 10.1590/1678-4324-2021200817

Michl J, Park KC, Swietach P. 2019. Evidence-based guidelines for controlling pH in mammalian live-cell culture systems. Communications Biology 2: 10.1038/s42003-019-0393-7. DOI: 10.1038/s42003-019-0393-7

Using Phenol Red to Assess pH in Tissue Culture Media. (2018). https://www.semanticscholar.org/paper/Using-Phenol-Red-to-Assess-pH-in-Tissue-Culture/53c1d3d3ffc060294c1d1e023e44a8eb97771e06

